# Visual Attention Through Uncertainty Minimization in Recurrent Generative Models

**DOI:** 10.1101/2020.02.14.948992

**Authors:** Kai Standvoss, Silvan C. Quax, Marcel A.J. van Gerven

**Affiliations:** Donders Institute for Brain, Cognition, and Behavior, Radboud University; Einstein Center for Neurosciences Berlin

## Abstract

Allocating visual attention through saccadic eye movements is a key ability of intelligent agents. Attention is both influenced through bottom-up stimulus properties as well as top-down task demands. The interaction of these two attention mechanisms is not yet fully understood. A parsimonious reconciliation posits that both processes serve the minimization of predictive uncertainty. We propose a recurrent generative neural network model that predicts a visual scene based on foveated glimpses. The model shifts its attention in order to minimize the uncertainty in its predictions. We show that the proposed model produces naturalistic eye movements focusing on informative stimulus regions. Introducing additional tasks modulates the saccade patterns towards task-relevant stimulus regions. The model’s saccade characteristics correspond well with previous experimental data in humans, providing evidence that uncertainty minimization could be a fundamental mechanisms for the allocation of visual attention.

## Introduction

Any organism interacting with a dynamically changing environment has to infer the causes of the observed dynamics through incomplete information from its sensory organs. It is bombarded with a continuous stream of sensations which are noisy and largely redundant to create a useful model of its surrounding. Such a model needs to faithfully capture the latent structure of the world, while being adaptive enough to allow for rapid interactions and updating. At the same time, the brain’s cognitive resources and processing capacities are limited (***Marois and Ivanoff, 2005***). Consequently, the ability to filter unimportant input and sequentially focus on interesting stimulus regions is a key mechanism of neural information processing. As such, attention plays a fundamental role in cognition at large, not only for processing of external afferents but also in internal processes like memory (***Chun and Turk-Browne, 2007***) and decision making (***Orquin and Loose, 2013***).

In human perception, vision is the most dominant sense and has evolved to efficiently process enormous amounts of information using only a limited-bandwidth information channel (***Itti and Koch, 2001***). A significant portion of the brain is dedicated to visual processing (***Van Essen et al., 1992***) and our eyes allow us to rapidly focus on different parts of the visual field (***Thorpe et al., 1996***). The eyes’ anatomy favors the selective processing of relevant stimuli through a central foveal region that processes information with high resolution and a periphery that has a lower resolution. Overt attention — moving an attended region of the visual field into the fovea for heightened processing — is a central selection mechanism to make sense of a visual scene. This active sampling of the environment requires quick saccadic eye movements that bring relevant parts of a scenery into focus. Given the relevance of the visual sense, eye movements and visual attention remain an active field of research (***Moore and Zirnsak, 2017***).

A question central to this research is how the brain allocates its attention and decides which locations to sample through saccades when observing a visual scene. As mentioned, attention is intricately linked to many cognitive processes and is thus not only affected by external stimulus features. Instead, endogenous variables like expectations (***Neider and Zelinsky, 2006***), emotions (***Kaspar and König, 2012***), and task requirements (***Land and Hayhoe, 2001***) play an equally important role. While the effects of bottom-up stimulus properties as well as top-down processes on visual attention have been studied extensively in isolation, fewer accounts have investigated their interaction and sought to explain both aspects within a comprehensive theoretical framework (***McMains and Kastner, 2011***).

The most prominent account of bottom-up processing holds that attention and saccades are primarily affected by the saliency of observed stimuli (***Itti et al., 1998***). In this view, several maps of different visual features like “color” and “orientation” are generated in parallel in early visual areas over the entire observed scene, and then combined into a single saliency map (***Koch and Ullman, 1987***). The most salient stimulus region, i.e., that part of the observed scene that “stands out” the most, determines where attention is directed to. Several computational models of this kind of bottom-up processing have been proposed (***Koch and Ullman, 1987***; ***Itti et al., 1998***; ***Le Meur et al., 2006***) and saliency models have predicted eye-movements well under some experimental conditions. However, they cannot account for all aspects of attention allocation (***Schütz et al., 2011***) and often only describe processing of basic features like color or orientation.

Most evidently, classical saliency-based models do not address top-down attentional influences like task requirements. The weighting of certain object features can be shifted according to task-dependent modulations and attention can be directed to entire objects instead of individual stimulus features (***Gilbert and Li, 2013***). While features vary across object boundaries, saliency-based attention alone cannot account for attentional shifts driven by object recognition (***Schütz et al., 2011***). This object-guided attention can occur very fast (***Thorpe et al., 1996***) and is a good predictor for saccadic eye-movements (***Einhäuser et al., 2008***). Since object recognition, and more general task demands that require planning of saccade paths, demonstratively affect saccade behavior above mere saliency, they are usually treated as separate components of attentional control (***Schütz et al., 2011***) and have been modelled independently (***Desimone and Duncan, 1995***; ***Oliva et al., 2003***; ***Reynolds and Heeger, 2009***). Nevertheless, both effects clearly interact in real-world scenarios in ways that are not yet well understood (***McMains and Kastner, 2011***).

A crucial insight for understanding the interaction between bottom-up and top-down processing is that attention allocation constitutes an *active* information seeking process (***Gottlieb et al., 2013***) rather than a mechanistic response to salient stimulus properties. In this view, attention serves to maximize the information gained from each saccade. From an information-theoretical perspective this corresponds to minimizing global uncertainty (***Guiasu and Shenitzer, 1985***). In free viewing, these regions of maximal uncertainty correspond to salient stimulus regions which, by definition, are harder to predict given their surrounding, while in a task setting, uncertainty is shaped by task-relevant prior expectations. The former addresses sensory uncertainty whereas the latter involves uncertainty about the latent causes of the visual percepts. From a predictive coding perspective, however, these two processing levels interact through a generative model that describes how the latent structure generates the observed sensations (***Huang and Rao, 2011***). Therefore, the idea that visual attention serves to resolve uncertainty about the observed visual scene parsimoniously conjoins both aspects of attention within a single objective. Similar formulations describe this process in terms of Bayesian surprise (***Baldi and Itti, 2010***) or free-energy minimization (***Feldman and Friston, 2010***).

Empirical evidence indicates that this generative framework is not merely an elegant explanation but also describes human performance better than other accounts (***Renninger et al., 2007***; ***Mirza et al., 2018***). However, ***Renninger et al.*** (***2007***) and ***Mirza et al.*** (***2018***) also state that heuristic formulations rather than Bayesian models of global uncertainty can describe the data better. This indicates that mathematical formalizations akin to the one offered by ***Mirza et al.*** (***2016***) seldomly capture the intricacies of complex neural computations, despite their biological accuracy. Moreover, the level of abstraction of these models often makes it difficult to interpret the fundamental neural components at play.

Instead, we propose an artificial neural network (ANN) model, that is loosely inspired by neural computation in the brain, in order to gain a deeper understanding of the role of uncertainty minimization in visual attention. Despite this loose coupling, ANN models have successfully revealed relevant computational principles of neural information processing (***Güçlü and van Gerven, 2015***) and it has been proposed that ANNs offer a valuable tool for exploratory research (***Cichy and Kaiser, 2019***). ***Kietzmann et al.*** (***2018***) argue that ANNs can help reveal mechanistic explanations of cognitive functions and drive insights in computational neuroscience by stripping away much of the biological detail of more faithful biophysical models.

ANN models have been proposed for saliency prediction (***Kümmerer et al., 2014***; ***Kroner et al., 2019***) as well as top-down modulation (***Stollenga et al., 2014***; ***Zhang et al., 2018***). But similar to more classical computer vision models like the ones proposed by ***Mnih et al.*** (***2014***) and ***Gregor et al.*** (***2015***), these models often do not take specific neuroscientific models into consideration, or treat bottom-up and top-down processing in isolation. Therefore, their potential to generate new hypothesis about the neural implementations of selective visual attention to inform experimental research are limited. A few other works have taken a more integrative approach and explicitly translated findings from computational neuroscience models into ANN architectures. ***Adeli and Zelinsky*** (***2018***) propose a deep neural network model that loosely mimics brain areas in the visual hierarchy and performs visual search by implementing biased-competition theory (***Desimone and Duncan, 1995***). Their model balances an adequate level of abstraction and biological plausibility but is computationally very expensive as it employs an entire grid of neural networks in order to mimic retinal vision with heightened foveal processing. ***Hazoglou and Hylton*** (***2018***) by contrast, do not explicate a specific neural circuit model but propose a simple ANN architecture that performs saccades on movie data through the minimization of hierarchical prediction errors. Their work is similar to ours in taking inspiration from predictive processing but uses only feed-forward neural networks and does not address the precision or uncertainty of the model’s predictions.

Works by ***Mirza et al.*** (***2016***) and ***Yang et al.*** (***2016***) demonstrate convincingly that integration of information over a sequence of saccades in order to minimize uncertainty explains visual attention allocation parsimoniously and at the same time predicts human behavior. In this vein, the goal of our research is to construct a ANN model to better understand the involved mechanisms of visual attention allocation. By using pixel values as direct input to the model and training it under different task requirements, both bottom-up as well as top-down processes are investigated. The proposed generative recurrent latent variable model is linked to computational attention models in the literature but exploits the benefits of ANN modeling. Thereby, the effectiveness of uncertainty minimization as a mechanism to learn relevant saccade paths is studied in detail. Specifically, our model enables the investigation of the interaction between top-down and bottom-up processes on a mechanistic level.

## Model

We propose a generative recurrent neural network model trained to perform a series of saccadic “eye movements”. The saccades constitute samples of a visual scene which the model needs to integrate in order to learn about the latent structure of its input. While each saccade reveals only a limited field of view, the model is optimized to make accurate predictions of the full visual scene. Trained as a variational autoencoder (VAE) (***Kingma et al., 2014***), the model learns a probabilistic latent representation and a generative model of the input data. This latent code is used to draw samples from the conditional posterior distribution given the sequence of prior saccades. Each of these samples describes a different prediction of the full input image given the available information.

The uncertainty in these predictions is used to guide the model’s subsequent saccade. Thereby, the model performs saccades in order to minimize predictive uncertainty.

Figure 1 depicts the network architecture. Information is processed in two streams as proposed by ***Mnih et al.*** (***2014***): a *where*-pathway which represents the currently attended location and a *what*-pathway which processes the foveated visual input. A retina-like visual field is simulated by downsampling consequtive image patches of increasing size to the resolution of a 4×4 px fovea. Hence, the model processes information with high acuity only at a small foveal region and perceives peripheral stimulation with decreasing resolution. Spatial coordinates are available to the network normalized to the range [−1, 1] with (0, 0) at the image center. Both pathways are instantiated by two-layer perceptrons.

**Figure 1.**
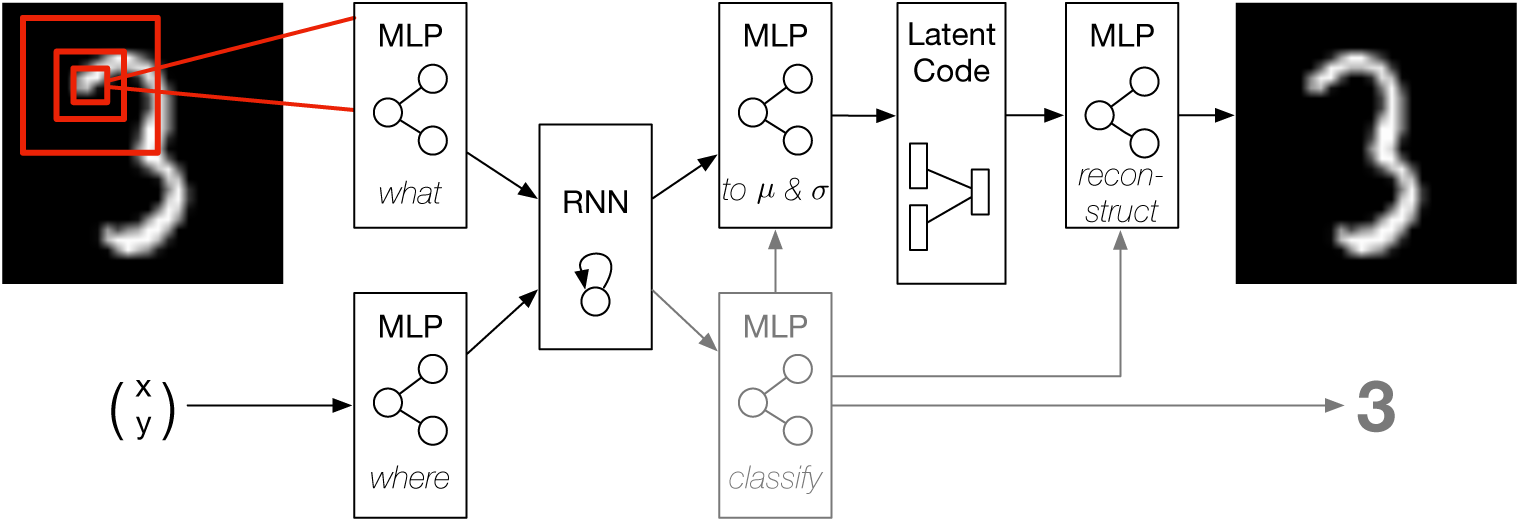
Network Architecture. The recurrent generative neural network architecture. Inputs to the network are the currently attended coordinates and the foveated glimpse. Feature representations of both are integrated in a recurrent layer. The recurrent representation serves to remember past experience and is used to parameterize a Gaussian latent representation. Samples from the latent code are used to generate predictions by the decoder. Variance in the predictions is used as a measure of pixelwise intrinsic uncertainty. In supervised settings, the recurrent representation is additionally used to train a classifier (gray). The classification output is used as input to the latent network and the decoder, disentangling the latent representation into class and style variables.

The transformed feature representations of the two processing streams are integrated in a recurrent neural network over the course of a saccade sequence. The recurrent activation is used to parameterize the mean and variance of a latent multivariate Gaussian distribution with standard normal prior. Sampling from this conditional posterior, using the reparameterization trick proposed by ***Kingma et al.*** (***2014***), a decoder neural network generates predictions of the full input stimulus. The decoder network as well as the parameterization network, that transforms the recurrent activation to the latent mean and variance, are instantiated by two-layer perceptrons.

The network can be trained in an unsupervised manner, with the objective to create accurate predictions while learning a structured latent representation, or in. a supervised manner by using additional label information. In the supervised setting the recurrent activation is additionally used as input to a two-layer perceptron that is trained to predict the class label. In this case, the predicted class label is used as additional information to the parameterization and decoder networks. Conditioning the generative model on the predicted class label allows the network to represent the visual style information disentangled from the stimulus class.

After each saccade, multiple samples of the visual scene are generated by the neural network. The pixel-wise variance in these predictions is taken as the intrinsic uncertainty of the model and used to guide the subsequent saccade. For that, the coordinates of the pixel with highest uncertainty are selected as the next saccade target.

After completion of a saccade sequence of predefined length, a loss in calculated and used to optimize the network parameters. The loss consists of a reconstruction term, driving the network to make accurate predictions, a Kullback-Leibler (KL) divergence between the latent representation and its prior distribution, enforcing a structured latent code, and, in the case of supervised training, a classification loss. The network was implemented in PyTorch (***Paszke et al., 2017***) and trained with stochastic gradient descent. A detailed description of the training procedure and network parameters can be found in the Methods section.

## Results

The neural network model was evaluated in a series of experiments, probing distinct aspects of bottom-up and top-down processing in visual attention. We used the well-known MNIST hand-written digit dataset (***LeCun, 1998***) to test task-independent and task-dependent visual attention. Furthermore, we made use of the “translated” and “cluttered” MNIST datasets (***Mnih et al., 2014***) to investigate visual search.

### Visual attention as uncertainty minimization

We first investigated whether the model learns to perform meaningful saccades with uncertainty minimization of the reconstructed images as optimization criterion without further supervision. As the model only ever sees a limited part of the image, it has to generate saccade paths that sample the stimulus sufficiently to accurately reconstruct the target image.

Figure 2A shows an example saccade trajectory for one test stimulus. The first row shows the target image and indicates the visual field of the network in red. The second row shows the actual input to the network. The downsampled periphery was upsampled to its original resolution in order to visualize the network input in the size of the target stimulus. It can be seen that the resolution of the periphery decreases rapidly and the network’s visual field does not cover the full extent of the image. Consequently, the model has to perform saccades that sample the full input effectively in order to be able to reconstruct the image.

**Figure 2.**
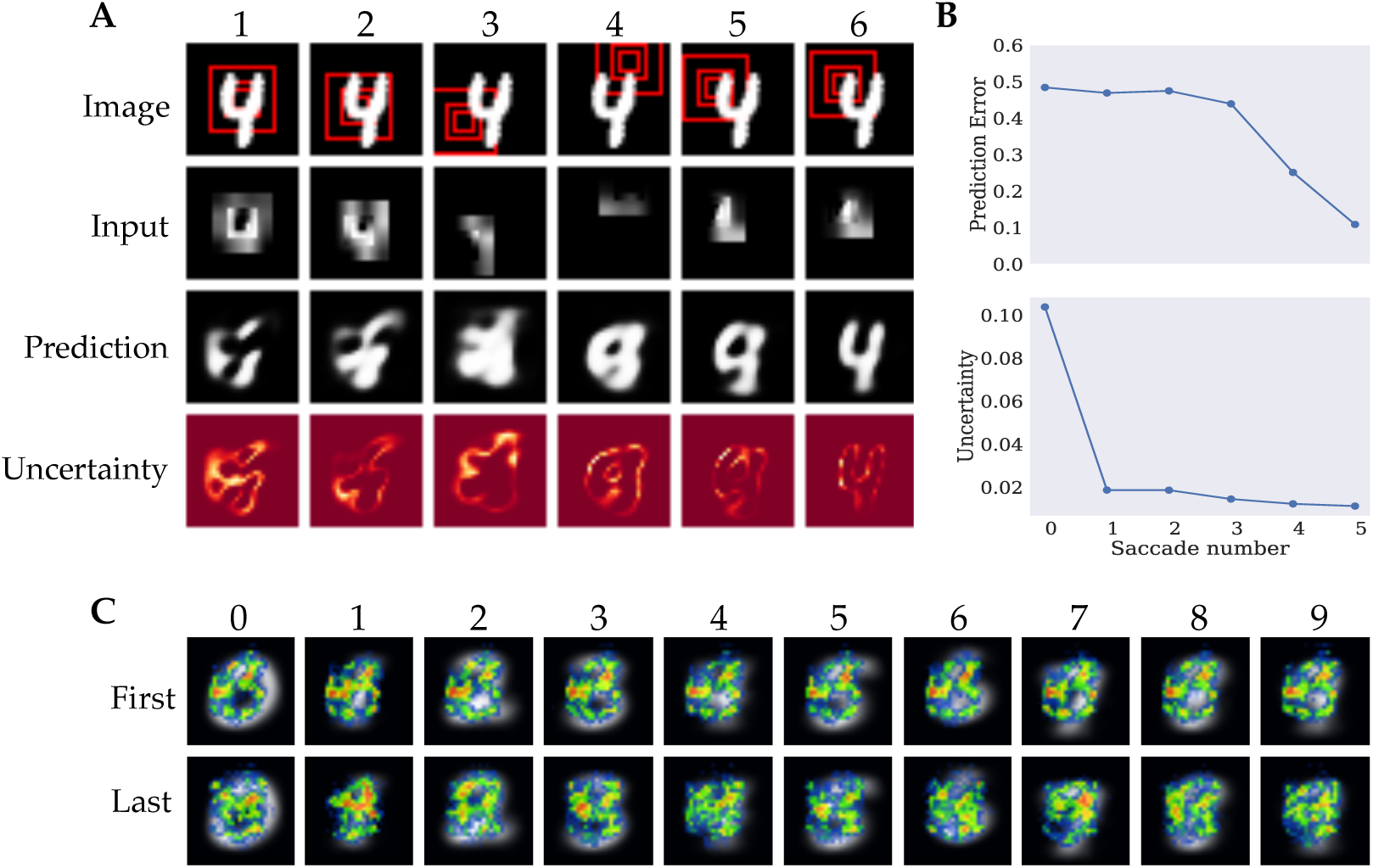
Unsupervised Model. A) Example episode of the model reconstructing a test stimulus with columns corresponding to individual fixations. The first row shows the target image and the currently attended location. Red squares represent different levels of resolution, with the smallest patch corresponding to the full resolution fovea and the larger patches to the lower resolution periphery. The second row displays the current input to the network. The third row shows the model’s prediction after receiving the current input and the fourth row shows the pixel-wise model uncertainty obtained through sampling with lighter regions corresponding to higher uncertainty. All images are normalized to the range (0, 1). B) Prediction error and model uncertainty unsupervised training. Upper plot shows the mean reconstruction error as measured by Binary Cross Entropy after each saccade. Lower plot shows the maximum uncertainty after each saccade. C) Heatmaps of saccade targets over the test set for unsupervised training. Heatmaps were generated separately for the first three (top) and last two (bottom) saccades in order to account for temporal patterns in the saccade paths. Saccade end-points were aggregated per digit over all test samples. Average digit representations are overlayed by the heatmaps for digits zero to nine from left to right.

To investigate whether the proposed architecture is able to integrate visual information over time and learn an adequate generative model of the image statistics, we visualized the model predictions after each saccade, in the third row of Figure 2A The depicted predictions show one individual sample of five forward passes that are used to quantify the intrinsic uncertainty. As the network samples the stimulus at different locations, the predictions become more accurate and replicate the target reasonably well at the final saccade. More examples of the network’s reconstruction after integrating a full saccade sequence can be seen in the appendix Figure 6. Additional videos of the network performing saccades can be found at https://github.com/kstandvoss/visual_attention. The reconstructions show that the model successfully learns to predict the images. The individual differences in the reconstructions from images from the same class show that the network does not merely generically reproduce the class identity but instead captures the respective style of the target.

Another relevant aspect of the model is its uncertainty estimation that is used to guide the saccade trajectories. The last row in Figure 2A shows the current model uncertainty after each saccade. More examples of uncertainty maps can be seen in appendix Figure 7. It can be seen that the model is most uncertain about digit contours which also correspond to regions of highest local contrast and thus salience and local uncertainty (***Renninger et al., 2007***). This can be best observed in the uncertainty predictions after the full saccade sequence as the image reconstructions best capture the target digits. In contrast, after the model has only observed the stimulus at the image center at the first saccade, the model predictions are still quite abstract and often do not resemble the class identity of the target. Consequently, the network uncertainty is more disperse and generally higher, resulting in a broader sampling of the digit.

This observation is further quantified in Figure 2B. The mean prediction or reconstruction error after each saccade, evaluated on the test data, is depicted on the top. The binary cross-entropy between predicted and target pixel values is used as a measure for the quality of the reconstruction. It shows that the model predictions remain relatively inaccurate for the initial saccades but steadily improve during the final glimpses. This implies that the model samples relevant information for predicting the target scene. The lower graph shows the uncertainty of the reconstructions summed over all pixels in the model prediction, averaged over the test data after each saccade. It shows that the uncertainty is reduced over the course of the saccade sequence. The maximum decrease in model uncertainty can be observed after the first saccade — while the prediction error only decreases slightly — and a further monotonic decrease in uncertainty can be seen for the final saccades.

The uncertainty estimation served the guidance of the model’s exploration of the input using saccadic eye-movements. Closer analysis of the saccade patterns can help us understand the policy that the model learned. One option is that the model just randomly samples the image without learning meaningful saccade patters. Another option is that a single strategy is learned that it applies to every image, without using the actual input information it receives. However, both these strategies would not be very interesting and do not resemble saccadic eye movement pattern humans typically show. Humans adapt their eye movement pattern to the particular object they are observing, using information from their previous saccades to guide the next one. Ideally, our model would show such eye movements. Heatmaps were visualized for each of the ten digit classes (Figure 2C). The heatmaps were generated by recording the target pixel location of each saccade separately for all samples in the test data, forming one density map of the network’s fixations for each of the *S* saccades. The density maps were then combined for the first three and last two saccades, respectively. The initial fixation of the image center was not included. Aggregating the density maps for the initial and final saccades separately keeps some notion of temporality while combining the data of several fixations whose patterns were very similar to each other. The similarity of the first saccade patterns and last two fixations could be related to the transition in reconstruction quality between the third and fourth saccade observed in Figure 2B. The first row of Figure 2C shows the combined density maps for the first three saccades per digit displayed over the mean image of the respective digit class. It can be seen that the sampling strategy during the initial saccades is very similar across all classes. In all cases it follows a rough three-shaped pattern in addition to a strong focus on the left center of the image. The second row shows the aggregated fixations for the final two saccades. Here, the saccade patterns differ more strongly from one another, indicating that the model learned to choose the next saccade location based on the observations during the preceding saccades.

### Analysis of fixation behavior

In order to compare the learned fixation behavior to behavior that humans typically show, it is useful to further analyze the saccade statistics. Figure 3A shows the distributions of saccade lengths — or amplitudes — as the euclidean distance between subsequent fixations in a normalized histogram (shown in blue). It can be seen that the histogram follows a Gamma distribution. To show that this is indeed the case, a truncated Gamma distribution was fitted to the data and visualized in orange (Γ(*α* = 11.90, *β* = 1.097)). The model almost never fixated locations further away than its peripheral field of view of 16 pixels. The mean saccade amplitude was 7.34 with a standard deviation of 3.74.

**Figure 3.**
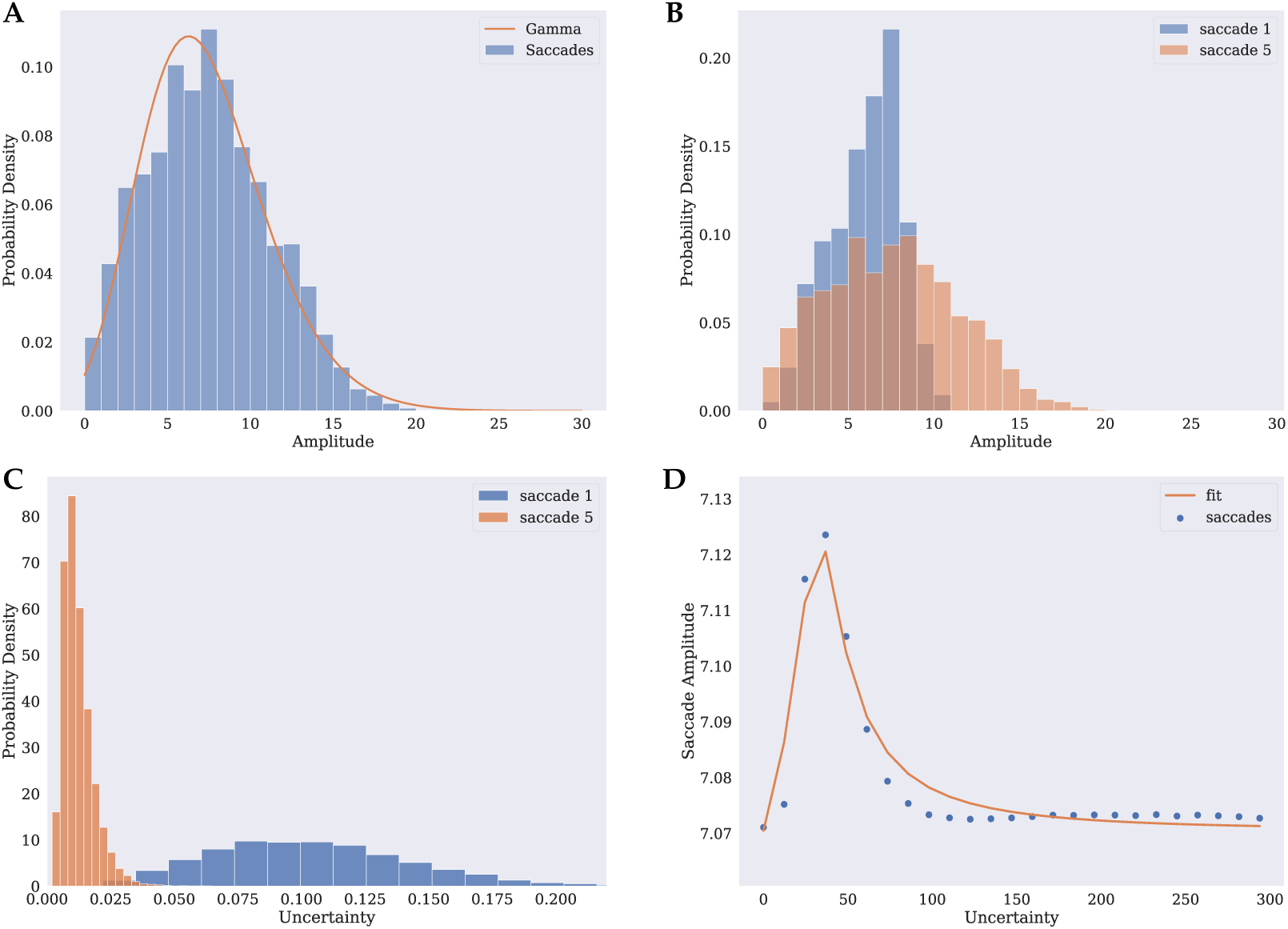
Overall Saccade Statistics. A) Histogram of saccade amplitudes (blue) and fitted Gamma distribution (orange). B) Histogram of saccade amplitudes for first (blue) and last (orange) saccade. C) Histogram over maximum uncertainty for first (blue) and last (orange) saccade. D) Saccade amplitude as a function of uncertainty as a proxy for fixation duration (blue) and fitted saccade model by ***Unema et al.*** (***2005***) (orange).

When separated into earlier and late saccades, different patterns become visible. While the first saccade is distributed very peaked at a mode at 7.07 with a slight right skew and a low mean of 5.83, the distribution of the last saccades is more left skewed with a mean of 7.41 and a wider range, including long saccades up to 20.81 pixels (Figure 3B). This difference in saccade amplitudes between earlier and later saccades matches the observation based on the heatmaps that the earlier saccades sample the digit more densely in the immediate periphery than the later saccades, which are distributed more evenly across the scene.

That human fixations follow a Gamma distribution has been reported in the literature for free viewing (***Ylitalo et al., 2016***) as well as for task conditions (***Castelhano et al., 2009***). Another variable that is usually investigated in human eye movements is the fixation duration, which also follows a Gamma distribution (***Castelhano et al., 2009***; ***Ylitalo et al., 2016***) and is correlated with the saccade amplitude (***Smit et al., 1987***; ***Unema et al., 2005***). Since our model does not feature a model independent temporal component, saccade durations cannot be measured. However, ***Koenig et al.*** (***2017***) showed that human fixation durations are prolonged with increased reward uncertainty in an associative learning task. If we generalize these findings, uncertainty could be taken as a proxy for fixation duration in our paradigm, following the intuition that regions of higher uncertainty should be sampled more accurately (***Henderson, 2017***).

Similar to the fixation duration reported in the literature, our model uncertainty follows a Gamma distribution, too (Figure 3C). When separated for the different saccades we can observe that the uncertainty is highest at the first saccade and roughly follows a normal distribution. At later saccades, the uncertainty decreases and is more left skewed towards zero. A similar behavior in human saccade latencies has been described by ***Loon et al.*** (***2002***), showing that the first saccade follows a more normal distribution with an increasing kurtosis towards later saccades.

In humans the relationship between fixation duration and amplitude is not linear, though, and instead shows a positive relationship for low durations and a negative relation for higher latencies ***Unema et al.*** (***2005***). Figure 3D shows the saccade amplitude as a function of uncertainty as a proxy for fixation duration in our simulation. Uncertainty was binned into evenly spaced bins. Data points represent the mean amplitude per bin. Additionally, a model of human saccade generation proposed by ***Unema et al.*** was fitted to our data, shown in orange. The uncertainty range has been rescaled to the range of human saccade latencies as reported by ***Unema et al.***. It can be seen that the model provides a good fit for the simulated data. The observed peak saccade amplitude for lower latencies can be observed similarly in human experiments.

### Task-dependent visual processing

In humans, the saccade patterns in response to an image can be strongly influenced by the task someone is performing (***Hayhoe and Ballard, 2005***). To investigate whether our model is able to modulate saccade patterns based on top-down task requirements, the model was trained in a supervised way to perform a classification task on the digits in addition to reconstructing the image.

Using model uncertainty to guide saccades, the model obtained an overall test error of 2.25% with a central initial fixation, or 3.59% when initialized at a random location at *t* = 0. In contrast, the same model performing random saccades obtained an error of 38.18%, highlighting that uncertainty driven saccades improved task performance.

To answer how the additional classification task affected the network saccades, the saccade patterns were analysed in more detail. Heatmaps of the supervised setting can be found in the supplementary material. Figure 4 shows an alternative visualization of the model’s saccade behaviour in terms of the mean pixel-wise uncertainty. Regions of high mean uncertainty are fixated more likely. Figure 4C and D show that the additional training objective affected the attention allocation. The first two rows of Figure 4C show the mean uncertainty after the first saccade for unsupervised and supervised training. It displays that both models preferably focus on the left center. However, the supervised model samples the image more evenly. Figure 4D depicts the mean uncertainty at the end of the saccade sequence for a selection of digits. It shows that at later saccades both models follow the digit contours more clearly. Yet, the supervised model samples relevant image regions more broadly. Despite comparable reconstruction performance, the supervised model uncertainty captures the digit shape more clearly. Another depiction of the more diverse saccade patterns of the supervised model can be seen in the appendix in Figures 9 and 10

**Figure 4.**
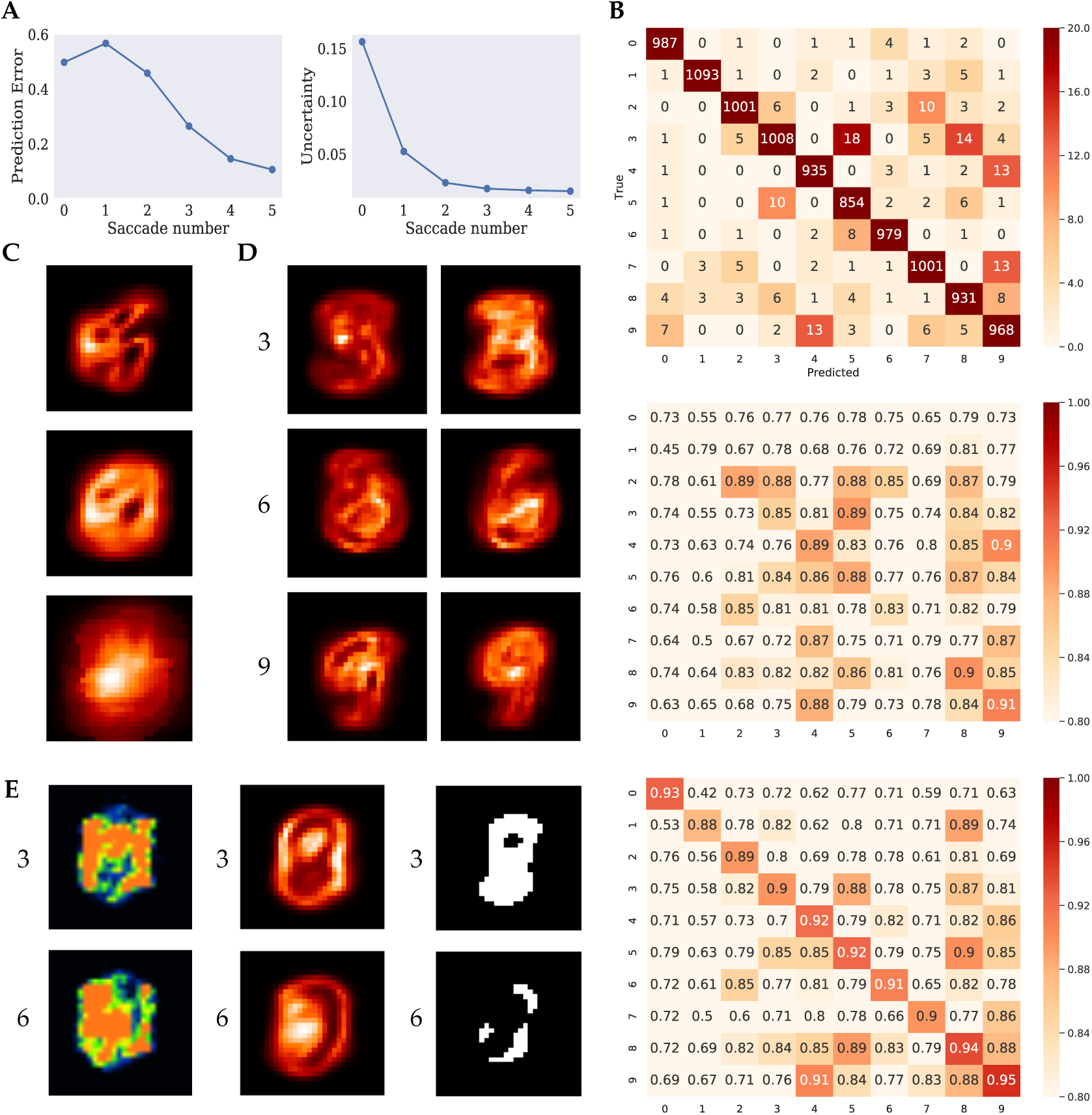
Model Comparison. A) Prediction error and model uncertainty for unsupervised training. Left plot shows the mean reconstruction error as measured by Binary Cross Entropy after each saccade. Right plot shows the maximum uncertainty after each saccade. B) Confusion Matrix (above) and correlation matrix between mean uncertainty maps and mean digits for unsupervised (center) and supervised (below). C) Mean pixel-wise uncertainty after first saccade for unsupervised (above) and supervised (center) training and pixel-wise classification score of logistic regression (below). D) Mean pixel-wise uncertainty after last saccade for the digits three, six, and nine, for unsupervised (left) and supervised (right). E) The first column shows the mean uncertainty for samples that were classified as belonging to a non-target class. The second column shows Heatmaps of saccade targets of binary task for all classes combined. The last column shows the thresholded mean of the most misclassified digit classes — {2, 5, 8} for threes and {0, 2, 4} for sixes — with high values shown in black.

One explanation for the difference in uncertainty estimation and consequently saccade patterns, could be that the additional classification criterion requires the network to commit to a digit classification more quickly and therefore produces earlier reconstructions of real digits. That this is indeed the case was further validated by the respective reconstruction error, which is displayed in Figure 4A. Compared to the unsupervised setup, the error decreases more rapidly. An effect of this could be that posterior samples for the supervised network are more likely drawn from a single digit category producing clearer uncertainty maps. Figure 4A also shows that the uncertainty in this experiment decreased slightly more slowly, yet also more smoothly than without supervision. The overall statistics of the saccade patterns as well as the uncertainty did not differ notably compared to the unsupervised training.

To better understand the observed differences in the initial saccade behaviour, a logistic regression classifier was trained on the raw pixel data of the full MNIST digits. Subsequently, for each pixel in the image a surrounding 8×8 patch was taken and the remaining pixels of the image were set to 0. This “foveated” image was used as input to the trained logistic regression classifier and the resulting score was taken as the value for the respective center pixel. In that way, a 30× 30 pixel map of classification scores was obtained, providing an estimate of the class information contained at each location given its immediate surrounding. Figure 4C shows the resulting scores in the bottom row. The classification map provides an explanation for the observed focus of both models on the center left image region. It can be seen that this region achieved the highest score and is thus most informative of the class identity. Interestingly, this indicates that the model trained in an unsupervised way, without knowledge of the class labels, sampled regions most relevant for determining the class identity. Nonetheless, clear differences between the two settings are visible. Correlating the variance maps with the score map of the logistic regression gives a correlation of *r* = 0.81 (*p* < 1*e* − 5) for unsupervised and a correlation of *r* = 0.90 (*p* < 1*e* − 5) for the supervised training. This shows that the classification model samples more regions that are relevant for classification than the model that is only trained to reconstruct the image.

To better understand the differences at the end of the saccade sequence despite similar reconstruction performances of both models, Figure 4B shows the confusion matrix of the supervised model. Investigating which stimuli where most confused by the model can help explain on which regions the model focuses its attention. The second and third matrix of Figure 4B show the mean pixel-wise uncertainty of the unsupervised and supervised network, respectively, correlated with the mean digit representation for each digit separately. High correlation values on the diagonal indicate that the model attends to the digit contours as the uncertainty map corresponds well to the digit appearance. The off-diagonal elements show that the network focuses on regions of digits that were most distracting. It can be seen that the supervised model features a much clearer diagonal than the unsupervised network, pertaining to the observation that the supervised uncertainty is better tuned to the digit style. Additionally, the off-diagonal of the supervised model more strongly resembles those of the confusion matrix. This indicates that model learns to focus its attention to discern the stimuli from other classes. In contrast, the unsupervised network focuses less on relevant stimulus regions for discerning digit categories.

While the described findings provide evidence that task requirements notably affected the model’s saccade behavior, the supervised model introduced an additional network component. Therefore, it is not possible to attribute the observed differences solely to the top-down task requirements. In order to further investigate in isolation how different tasks modulate gazing behavior for models employing the *same* architecture, we tested the supervised model on two structurally identical binary classification tasks. These tasks required the model to decide whether the observed stimulus corresponds to a target class. One network was trained to differentiate threes from all other digits and the second one to decide between sixes and non-sixes. In that, both models were identical to each other except for the target category. Both tasks were learned efficiently with accuracies of 99.64% and 99.26% for sixes and threes, respectively.

In the previous experiment, the different digit categories were not equally well classified. While the network was able to classify the digit 6 near perfectly, its performance on 3s and 9s was considerably worse. Figure 4B shows that the network misclassified 3s for other digits baring some visual resemblance. Specifically, the network frequently classified 3s as 5s or 8s. Similarly, the network often confused 9s and 4s and 7s. In order to optimally solve the binary classification task, the saccade patterns should be adjusted to the task in order to minimize these misclassifications. Therefore, we investigated whether the saccade strategy was affected by the task and whether the saccade patterns served the identification of the target class.

The first two columns of Figure 4E display the mean uncertainty and the saccade targets for the two task conditions. Both models focus on different regions of the input. The saccades feature very distinct sampling patterns that are visibly different from the saccade strategies seen previously. The network trained to distinguish threes focuses on the left and right sides as well as the upper half of the image. The network trained to classify sixes, in contrast, focuses more strongly on the left and lower left center. In order to gain a better understanding into what drove the saccades for non-target stimuli, the third column shows mean pixel-values taken over the classes that were most confused for the respective target class, thresholded to produce a binary relevance map. The confunders were the classes 2, 5, and 8 for the target class 3 and 0, 2, and 4 for the target class 6. It can be seen that the uncertainty — and the resulting fixations — corresponds well to the surrounding of the most confused classes, indicating that the network focused on those regions that helped discerning the target classes from the confounding digits.

### Visual search

The previous experiments focused on sampling of individual stimuli at the center of the image. However, a primary function of saccadic eye movements and attention in general is to find relevant information and select important input amongst distracting stimuli. For that reason, visual search tasks are often used to study eye movements and attention in humans. Here we investigate whether the proposed attention mechanism can be used to locate relevant stimuli within a larger scene and to discern them from irrelevant distractors. For that, we tested the model’s behavior on two visual search tasks. The first task was to find and classify an MNIST digit randomly located within a larger image without distracting stimuli. The second task was to find and classify the digit amongst distractors that consisted of cropped parts of other MNIST digits.

Figure 5 A shows exemplary search episodes for the translated MNIST task and Figure 5 B for the cluttered MNIST task. This model’s visual field was increased, shown in the figure’s first two rows, allowing it to process a larger extent of the input. The third row shows that the model predictions are considerably worse than in the previous experiments. As the digits are not centered, the model cannot know their location in advance. This uncertainty in the digits position is most visible after the first fixation. In both examples, the first glimpse does not reveal the full digit, so that the model mostly relies on an uniformed prior and produces very broad and blurred predictions. This behavior is more pronounced in the case of the cluttered data, since the information that is available to the model is not enough to determine the target location among the distractors. On the translated task, in contrast, the model does not have to deal with distracting stimuli and can therefore delimit the digit’s location more precisely. The model uncertainty about the locations is shown in the last row. In both cases, the models use the subsequent saccades in order to refine the prediction about the target location. In neither case does the model immediately fixate the digit but performs intermediate saccades that further inform it about the digit position. The model uses the final saccades to sample the digit after locating it. In the cluttered case, the model uncertainty reveals that the distractors cause the model to be less determinate about the position of the digit during the first two saccades than in the translated case.

**Figure 5.**
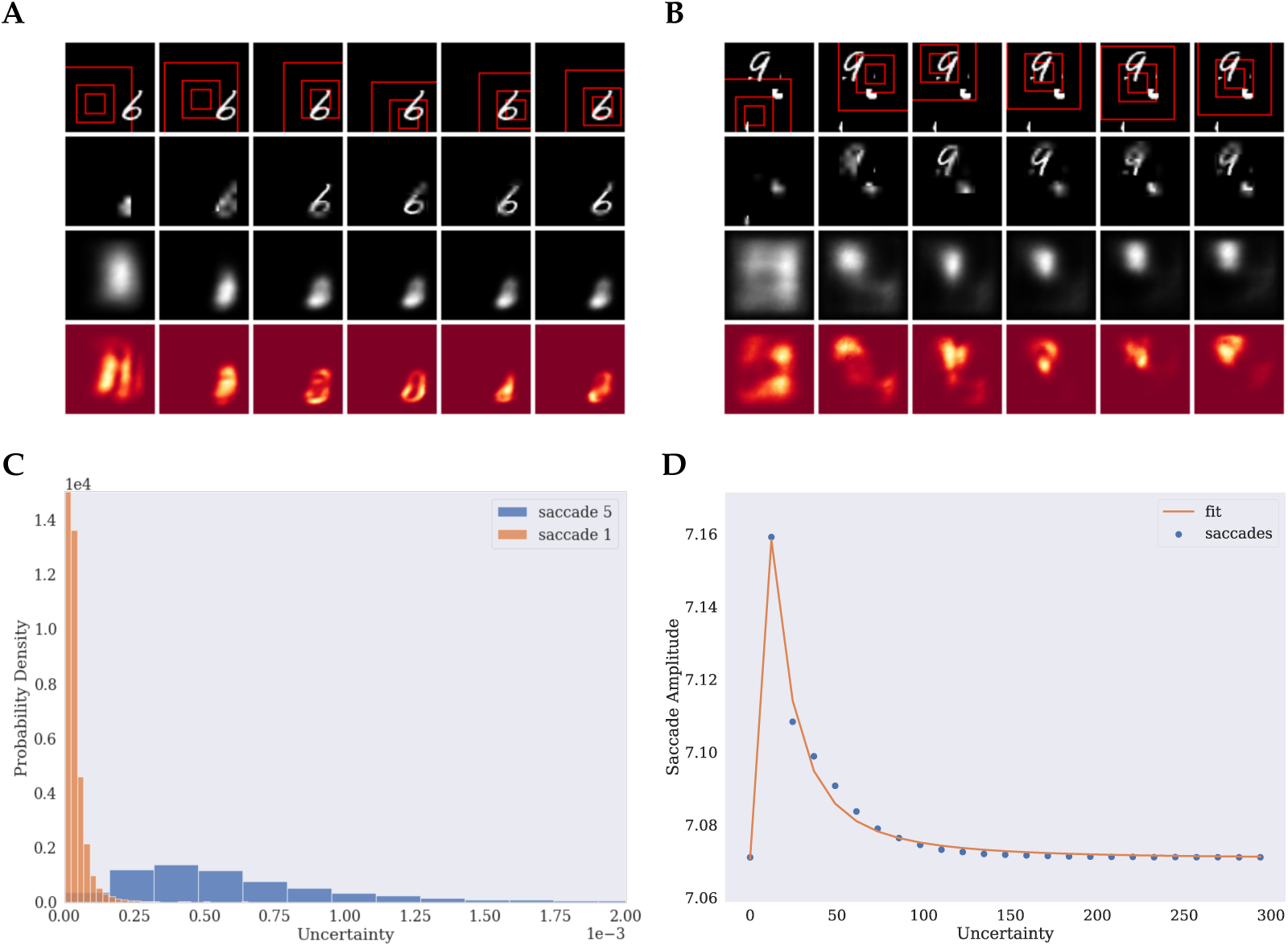
Visual Search. Example episode for the translated MNIST data (A) and cluttered data (B). C) Histogram over maximum uncertainty for first (blue) and last (orange) saccade. D) Saccade amplitude as a function of uncertainty as a proxy for fixation duration (blue) and fitted saccade model by ***Unema et al.*** (***2005***) (orange).

**Appendix 0 Figure 6.**
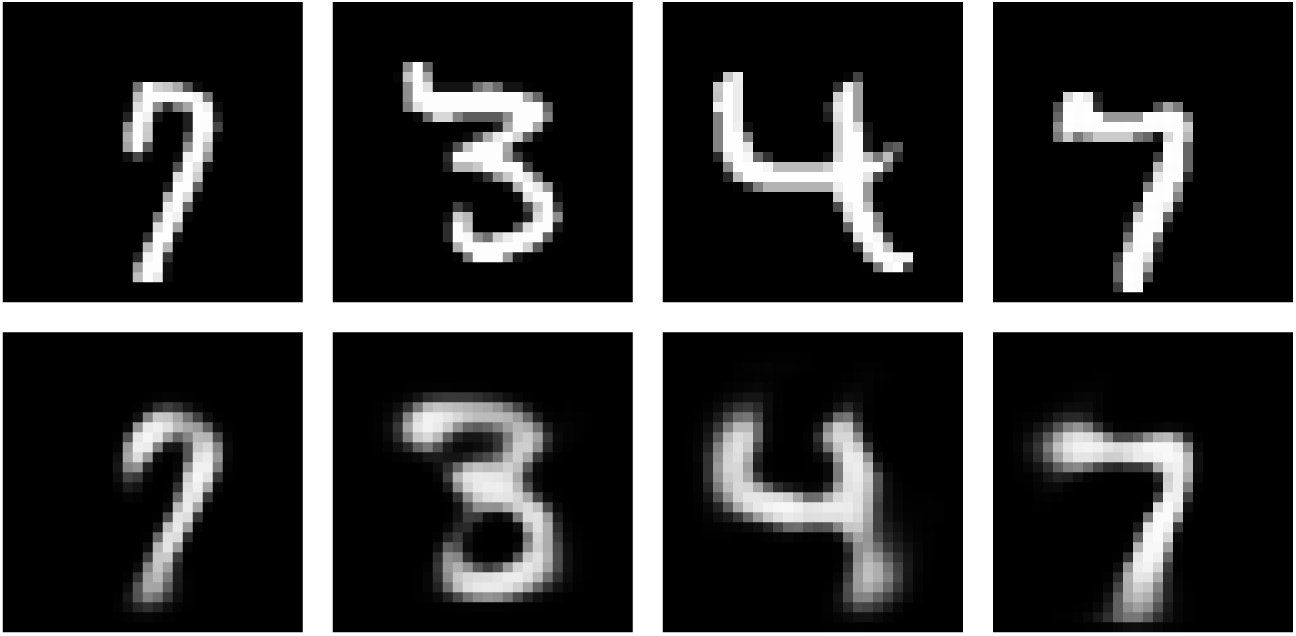
Example predictions after the final saccade. First rows shows the target stimulus, second row the reconstructed images.

**Appendix 0 Figure 7.**
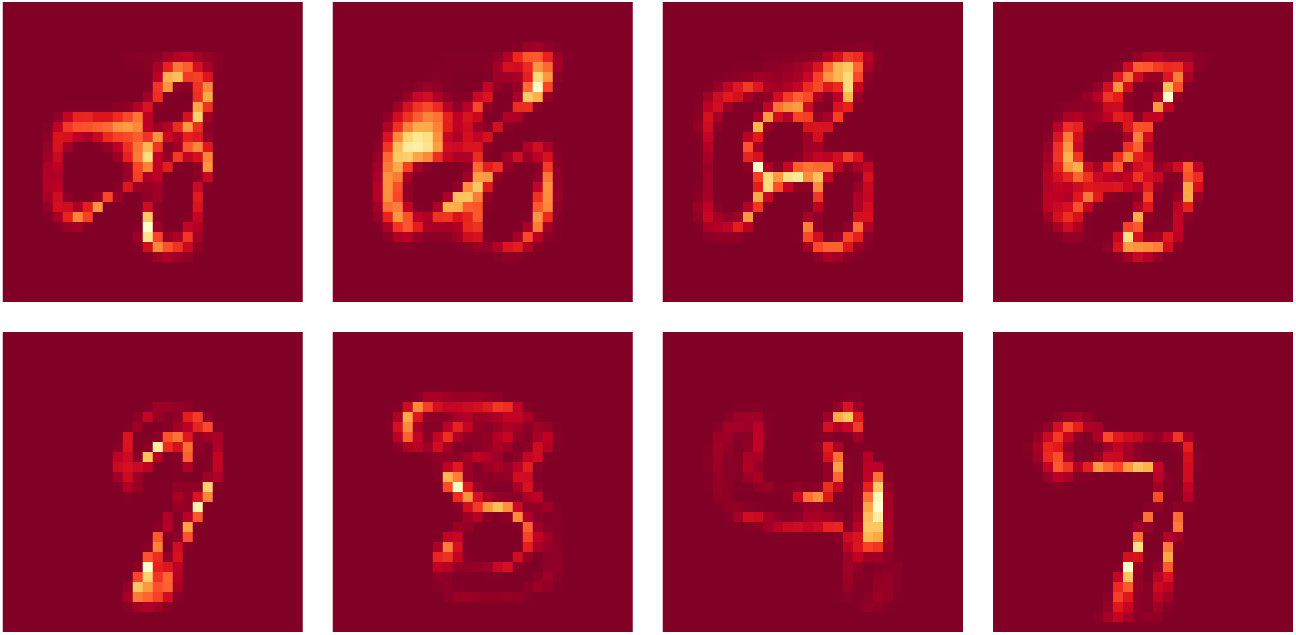
Example uncertainties after initial and final saccade. First rows shows the uncertainties after the first glimpse, second row at the end of the saccade sequence.

**Appendix 0 Figure 8.**
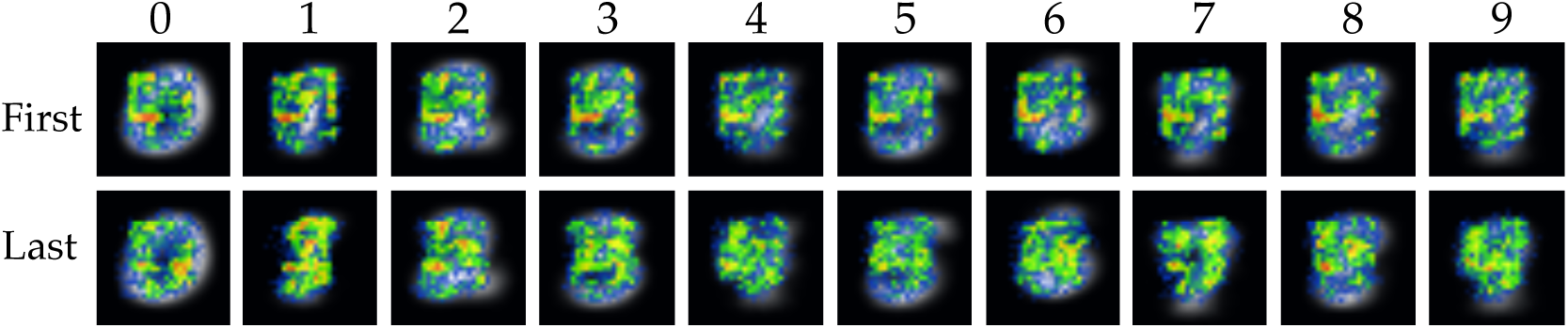
Heatmaps of saccade targets over the test set for supervised training. Heatmaps were generated separately for the first three (top) and last two (bottom) saccades in order to account for temporal patterns in the saccade paths. Saccade end-points were aggregated per digit over all test samples. Average digit representations are overlayed by the heatmaps for digits zero to nine from left to right.

**Appendix 0 Figure 9.**
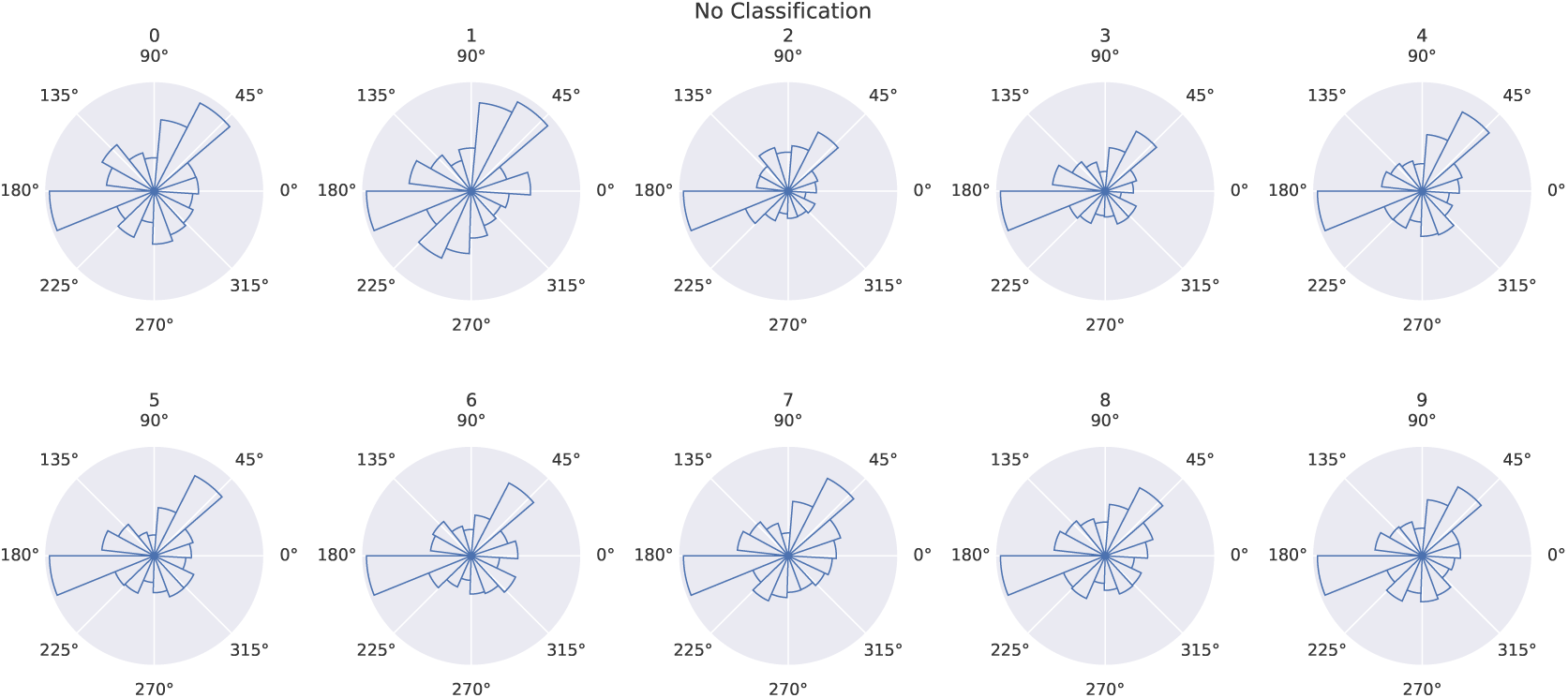
Circular histograms of saccade target locations per digit for unsupervised training. Bar height represents count of saccades towards the respective direction from the center. The similarity of all histograms reveals similar saccade behavior for all digits.

**Appendix 0 Figure 10.**
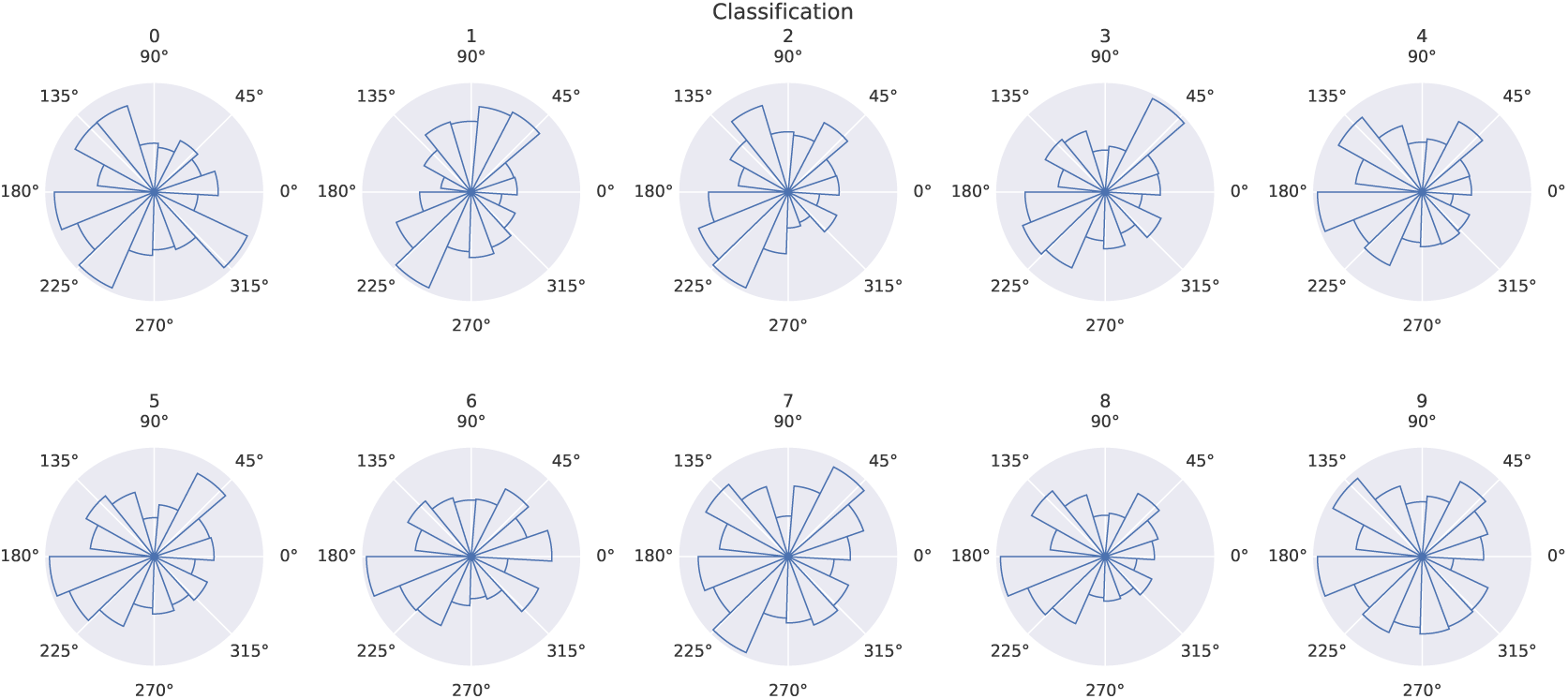
Circular histograms of saccade target locations per digit for unsupervised training. Bar height represents count of saccades towards the respective direction from the center. The model displays more diverse patterns than the unsupervised model.

The overall saccade statistics were similar to the previous experiments for both models. One difference can be observed in the mean saccade length per fixation. While the mean saccade lengths increased over time in the previous experiments, here, the saccades become increasingly shorter as the model finds and fixates the digit. This pattern reversal in contrast to the earlier experiments is visualized in Figure 5C. Figure 5D shows that the relationship between model uncertainty and saccade amplitude is again captured by the model proposed by ***Unema et al.*** (***2005***). In contrast to the other setting, the model displays a a sharper peak at low uncertainties.

That the learned search strategies are useful for classification is revealed by the obtained classification accuracies. Table 1 shows the classification errors for both datasets for the model with uncertainty guided attention as well as for random saccades. It can be seen that also in this case, uncertainty minimization is a useful attention allocation mechanisms for object classification in a visual search task. The random strategy still allows the model to classify the majority of test samples correctly but is considerably worse than the informed saccade strategy. Interestingly, the uncertainty guided model achieved an even lower error rate on the translated data than the model trained in the previous experiment. This shows that the increased fovea of the model facilitates the integration of the relevant class information and the ultimate classification.

**Table 1.** Classification Performance. Classification errors in percent for different initialisation and sampling strategies on the three different datasets.

|  | Data |  |  |  |
| --- | --- | --- | --- | --- |
| Strategy |  | Centered | Translated | Cluttered |
| Central init. |  | 2.25% | - | - |
| Random init. |  | 3.59% | 1.92% | 5.92% |
| Random fixation |  | 38.18% | 13.53% | 21.87% |

## Discussion

Attention is a fundamental mechanism of neural information processing and focusing on relevant stimulus regions through saccadic eye movements is a key ability of humans to interact within a dynamically changing environment. While both bottom-up influences on attention allocation as well as top-down task modulation are well studied, their interaction is not yet fully understood (***McMains and Kastner, 2011***). A parsimonious reconciliation of these two processes posits that both serve the common objective to minimize uncertainty about the sensed world (***Feldman and Friston, 2010***). Exploring this hypothesis, we proposed an artificial neural network model that makes predictions of an observed scene and performs saccades in order to minimize uncertainty in its predictions. Uncertainty in this model is represented implicitly by sampling from a generative model of the observed scenes. As different task requirements affect the model’s latent representations, the model uncertainty is affected both by bottom-up saliency features of the observations as well as top-down task demands. Uncertainty minimization was implemented heuristically by choosing the pixel with the highest pixel variance as the next saccade target.

A series of experiments was performed in order to investigate whether uncertainty minimization constitutes an effective attention mechanism for scene prediction, object recognition, and visual search. The different experimental setups enabled us to analyze bottom-up and top-down influences on the model’s saccade patterns. They revealed that the proposed sampling strategy allowed the model to perform useful saccades for the integration of task relevant information with and without explicit supervision.

### Model Performance

The first experiment showed that scene reconstruction, constrained by a prior on the learned generative model, can be used as an optimization criterion to train an ANN to effectively sample a larger scene and to successfully predict the full image based on foveated glimpses. The model integrated the information over the saccade sequence to gradually improve its scene predictions and to minimize uncertainty. The predictions during the initial fixations were inaccurate and similar for all digit classes and improved to more accurately resemble the target stimulus only during the later saccades. Next to the limited information that is available to the model at the beginning of the saccade sequence, another explanation for this behavior is that the reconstruction loss was calculated at the end of the sequence. Therefore, the early reconstructions were not explicitly trained to be accurate. By calculating the loss after each saccade the model could be constrained to make more accurate predictions early on. However, this is not only computationally more expensive, constraining the model’s earlier reconstructions could also unnecessarily limit the model from affecting its own sampling strategy. As fixation targets were determined by the model uncertainty and uncertainty was obtained by the variance in the model’s predictions, not constraining the early predictions gives the model more flexibility in learning a sampling strategy relevant for the task. In contrast, calculating the loss after each saccade forces the model to commit to a perceptual interpretation early on with only limited available information, potentially leading to suboptimal sampling of the input.

The observed uncertainty patterns showed that the prediction varied most strongly at the contours and edges of the predictions as well as faint regions. There are several explanations for this observation. One the one hand, the behavior can be explained methodologically, as samples from the approximate posterior *p*(*z* | *x*) are expected to be similar for the same input when the latent distribution is well captured by the model. Therefore, the samples should vary most in the details as opposed to featuring vastly different predictions of the data distribution. Furthermore, since the reconstruction loss is averaged over samples, producing very different predictions is punished. On the other hand, an additional part of the explanation, pertaining to the image data itself, is that edges and regions of high contrast are also regions of high salience and local stimulus uncertainty (***Renninger et al., 2007***). Therefore, it it would be a useful and biologically plausible sampling strategy, under the proposed attention model, to focus on edges in the observed image.

It was seen that the model focused on these regions specifically during later saccades. It was also only during these later saccades that the uncertainty was found to be well calibrated by matching regions of high prediction error. Consequently, modeling attention as uncertainty- or prediction error minimization as proposed by ***Hazoglou and Hylton*** (***2018***) might display similar behaviours at least for later saccades.

The second experiment revealed that the proposed attention mechanism can also successfully be employed in a supervised task setting. The implicitly learned saccade strategy enabled our model to achieve comparable classification performance to similar models in the computer vision literature (***Mnih et al., 2014***) without the requirement to train the saccade strategy using reinforcement learning. The lower accuracy can be explained by the fact that a few digits were often misclassified for digits that were visually similar. Consequently, our model did not sufficiently sample regions that were necessary to distinguish these difficult cases. A potential explanation for that could be that the model’s objective requires a certain trade-off between classification accuracy and reconstruction. While this trade-off might have prohibited the model to achieve better classification scores, it is this interaction between task-requirements and stimulus features that we were interested in. Another reason for the observed difference could be that we used a smaller foveal region with a larger periphery than ***Mnih et al.*** (***2014***), since we were interested in investigating the influence of retina-like vision. This could have impeded the differentiation of certain digits. Evidence for this explanation is the higher classification accuracy obtained in the visual search task during which the model had a larger fovea to process the digit.

This visual search paradigm is more frequently used to study human saccade behavior. We showed that the uncertainty minimizing sampling strategy helped to find the relevant stimulus, even in the presence of distracting stimuli, while achieving competitive classification scores to ***Mnih et al.*** (***2014***). However, in contrast to the previous experiments the model did not learn to accurately reconstruct the target images but produced rather blurry images. This can be explained by the fact that the model trained on a pixel-wise loss. Since the digits are not centered, the model receives a large loss when it predicts the target location slightly shifted. Therefore, it is a better strategy to make blurred predictions to minimize the loss. A potential way of solving this issue could be to use perceptual losses (***Goodfellow et al., 2014***) instead of the pixel-wise binary cross-entropy.

### Task Dependence

One main research question we were interested in was how different task demands affect the selection of saccade targets. We observed that the introduction of a classification task did visibly affected the saccade patterns. An analysis of the pixel statistics revealed that the supervised model did focus more strongly on stimulus regions relevant for classification than the model trained only on scene prediction. A comparison with a model performing random saccades showed that the proposed sampling strategy achieved considerably better classification results. Hence, the models clearly adjusted the saccade strategy according to the respective task requirements. This was also seen in the fact that the supervised model focused more on regions relevant to distinguish the digits from their confounding classes than the unsupervised network.

To control for the difference in architecture and to isolate the top-down influences, the binary classification task further explained the effect of task requirements on the saccade patterns. Here, analysis of the saccade patterns revealed again that the model focused on those regions that were relevant to distinguish the target class from the most confounding classes without explicit optimization to do so. This highlights once more that the proposed attentional mechanism successfully conjoins bottom-up and top-down processing with a single objective.

### Saccade Behaviour

Further analysis of the model’s saccade patterns revealed several characteristics of human eye movements. It was shown that the model’s saccade amplitudes followed a Gamma distribution which has also been shown in the human visual attention literature (***Ylitalo et al., 2016***; ***Castelhano et al., 2009***). The simulated initial saccades were shorter (*μ* = 5.83) than the final saccades (*μ* = 7.85) following a similar linear trend as reported by ***Castelhano et al. (2009)*** for visual search and memorization tasks in human participants. However, other studies like the work by ***Antes*** (***1974***) have reported an opposite behavior with initial saccade amplitudes being larger, progressively decreasing for later saccades. ***Antes*** (***1974***) motivate their findings by arguing that initial saccades sample the observed scene more broadly and concentrate on informative regions while later fixations focus on image details. ***Castelhano et al.*** (***2009***) argue that their deviating findings might be a result of their scene memorization task which requires subjects to focus on details from the beginning of the trial. This interpretation would be in line with our findings, as our model is initialized at the image center and can thus similarly focus on stimulus details from the start, without the need of broadly scanning the scene. Our findings additionally suggest that later saccades serve the disambiguation of similar stimuli and thus feature both large range saccades to relevant stimulus regions as well as shorter, more detailed saccades as evidenced in the broader amplitude distribution. This is in correspondence with the suggestion by ***Antes*** (***1974***) and ***Brockmann and Geisel*** (***2000***) that eye movements sample informative scene regions and try to cover these regions most efficiently. Here, informativeness is expressed in terms of model uncertainty. The uncertainty distributions per fixation followed a Gamma distribution of more Gaussian shape for the first saccade and increasing left skew for later saccades as uncertainty was minimized. The close resemblance of these statistics to the development of human fixation durations (***Loon et al., 2002***) has lead us to the hypothesis that uncertainty, and thus informativeness, might be a good approximation for the time spend processing the information at a sampled location, which has been proposed before (***Antes, 1974***; ***Henderson, 2017***). Under this assumption we could capture the non-linear relationship between saccade amplitude and duration observed in humans (***Unema et al., 2005***) surprisingly well. ***Unema et al.*** (***2005***) seek to explain the observed relationship by the interaction of two processing systems operating at different time scales. One faster saliency processing system, and a slower inhibitory top-down process. Our model offers the interpretation that these systems are in fact realized by the same model that accumulates information over time. Therein, for long fixation durations and thus high uncertainty, the model has not yet formed a strong prediction of the latent stimulus structure and therefore samples the scene locally to form a hypothesis. With increasing evidence accumulation and thus lower uncertainty the model focuses on disambiguating stimulus regions as described previously. Finally, once the uncertainty approaches zero the need for sampling the scene decreases. Further comparison of our model to human fixations on the same image data could help a better understanding of these mechanisms. Multiple studies have reported an increase in fixation duration especially during the first saccades (***Antes, 1974***; ***Unema et al., 2005***; ***Castelhano et al., 2009***), whereas uncertainty in our simulations decreased over time in the first experiments. However, our model was initialized at the image center and thus did not require short latency exploratory saccades in order to find informative stimulus regions. As such, these findings can be explained similarly to ***Castelhano et al.*** (***2009***) for the short saccades they observed. This is in line with the pattern reversal found in the visual search task in which uncertainty is higher for later saccades. Therefore, our model predicts an increase in fixation durations for visual search tasks and a decrease for object recognition tasks involving single centered objects in human experiments.

The saccade patterns in the binary classification task indicated that the initial saccades were mostly driven by the top-down task requirements compared to later saccades. While it appears unintuitive that top-down processes have such an early effect, ***Reingold and Glaholt*** (***2014***) showed that target-distractor similarity can have an effect on human saccade behavior as early as 26ms after fixation onset. Moreover, the network was heavily trained to perform the task and could therefore develop sampling strategies prior to the start of a trial. Hence, our results provide further evidence against the popular view that visual attention is influenced by a slow acting top-down system and quick bottom-up processing and instead is heavily guided by top-down interactions throughout (***Chen and Zelinsky, 2006***).

## Conclusion

Altogether, the results showed that minimizing predictive uncertainty is a useful attention mechanism to effectively sample a stimulus, that can account both for bottom-up as well as top-down influences. Our findings add additional empirical evidence to the hypothesis that both attentional processes serve the same objective of focusing on informative stimulus regions, as they integrate well with the existing literature on visual attention. More importantly, we showed that neural network models offer a valuable tool to test existing hypotheses and explore new research ideas. Our model offers new insights into conflicting findings in the literature and future research into the mechanisms of our model can potentially shed new light onto the neural underpinnings of visual attention. To that end, the model should be tested on naturalistic datasets to see whether the model can hold up to more realistic scenarios. An interesting avenue to pursue is to investigate the differentiation of uncertainty in aleatoric and epistemic uncertainty (***Kendall and Gal, 2017***), as it is important for the model to be able to judge what uncertainties constitute irreducible noise in the data, which it should ignore, and what is resolvable uncertainty that should be attended. In that way, the model might be able to make more accurate saccades in the cluttered dataset and learn realistic eye movements in natural datasets. Furthermore, more extensive study of the learned latent representations as well as future investigations of more expressive generative models and alternative uncertainty metrics could generate new insights into the mechanisms of neural uncertainty estimation. An extended model trained on natural data holds a lot of potential when compared to human data for the same task and might even be used to reveal important variables to characterize non-pathological and pathological attention behavior (***Mirza et al., 2016***), e.g. by investigating the relevance of specific model components like the knowledge of the own fixation location (***Thakkar and Rolfs, 2019***). To conclude, the neural network model that we proposed provides a new approach to investigate the mechanisms that may underlie human visual attention.

## Methods

### Data

In all experiments the MNIST digit dataset (***LeCun, 1998***) was used, consisting of 60000 images of handwritten digits from 0 to 9 of size 30 × 30 pixels. In the visual search tasks, MNIST digits were embedded in a larger black background of 60× 60 pixels at random locations, resulting in a “translated” MNIST dataset. A “cluttered” dataset was created by additionally adding distractor stimuli of cropped digit parts to the translated data as proposed by ***Mnih et al.*** (***2014***).

The model received only a limited view of the full input images. *Hard attention* was implemented by cropping a foveal part of the scene around the attended location and peripheral patches of increasingly lower resolution. For the MNIST data, foveal patches were of size 4×4 pixels. Peripheral patches of 8×8 and 16 × 16 pixels were downsampled to the resolution of the fovea through bilinear interpolation. In the visual search experiments, patches were of size 12 × 12, 24 × 24, and 48 × 48. All patches were flattened and concatenated as input to the network.

### Model Architecture

The model consisted of a latent variable encoder-decoder architecture that was trained as a variational autoencoder (VAE) (***Kingma and Welling, 2013***) with the objective to reconstruct the full visual scene based on the foveated *glimpses*.

The encoder’s what- and where-pathway were instantiated as 2-layer MLPs with leaky rectified linear (ReLU) activation functions (***Maas et al., 2013***), whose output was concatenated and combined in another single layer of leaky ReLU units. This combined feature representation of image and location information was integrated over time in a recurrent layer with hidden activation

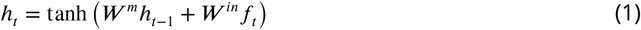

where *W ^m^* and *W ^in^* correspond to weight matrices that multiply the previous hidden state *h*_*t*−1_ and the new feature representation *f*_*t*_ respectively. The output of the recurrent layer was used to instantiate the approximate variational posterior distribution of the generative model *p*(*z* | *x*) with standard normal prior *p*(*z*) ∼𝒩 (1, 0). For that, the hidden representation *h*_*t*_ was used to parameterize the mean and variance of the posterior *p*(*z* | *x*)= 𝒩 (*z* | *μ, σ*) with

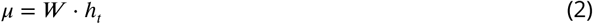

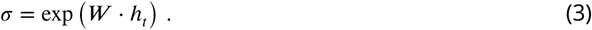

Since the latent code itself was not recurrent and only the current hidden representation of the recurrent layer is used to instantiate the mean and variance of the latent code, the recurrent layer has to integrate all relevant statistics of the image *x* in order to accurately capture the posterior distribution.

The *decoder* received samples *z* from *p*_*enc*_ (*z* | *x*) to generate the full image as the mean of a multivariate Bernoulli distribution. Samples from the posterior were drawn by sampling from the standard normal prior and combining the sample with the predicted posterior mean and variance. The mean of *p*(*x* | *z*), i.e. the reconstructed image, was computed from *z* with a 2-layer MLP. The first layer consisted of leaky ReLU units and layer normalization (***Ba et al., 2016***), and the final layer of sigmoid units.

In order to obtain an uncertainty estimate, *S* = 5 samples were drawn from the posterior and used to reconstruct the image. The variance from these predictions was taken as the model uncertainty

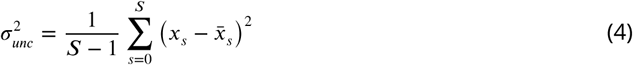

with forward predictions *x*_*s*_ and mean prediction 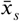. This pixel-wise model uncertainty was used to guide the subsequent saccade, with target location

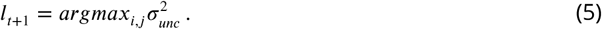

Subsequently, the newly attended location *l*_*t*+1_ was used to generate the foveated input to the network for the next time step. In that way, the model performed a total of *T* = 6 saccades, including the initial fixation, after which the loss was calculated and backpropagated through the network. The model was trained to minimize the binary-crossentropy (BCE) reconstruction loss, i.e. the log likelihood of the generative Bernoulli distribution, and the Kullback-Leibler (KL) divergence between the approximate posterior and the prior distribution

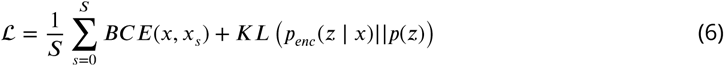

The reconstruction loss was averaged over all of the final samples *x*_*s*_ from the decoder. The loss term did not include the model’s uncertainty, hence uncertainty is only implicitly reduced by sampling locations of maximal uncertainty.

The network was trained using stochastic gradient descent (SGD) with nesterov momentum and weight decay. The learning rate was linearly decreased over the course of the training. The model was trained until the validation loss plateaued for 150 epochs or a maximum of *N* = 1000 epochs was reached. The validation loss was calculated after every epoch on a random held out subset of 10% of the original dataset. All MLP layers consisted of 128 units, the recurrent layer of 512 units, and the latent code of 25 variables. The KL loss-term was gradually tuned in using a weight parameter *β* as proposed as “deterministic warm-up” by ***Sønderby et al.*** (***2016***), but instead of using a linear scaling of *β*, it was determined by *β* = tanh(*ω* · *epoch*) where *ω* is a hyperparameter of the training procedure. It was set to 0.01 for all experiments.

In the supervised setting, additionally at each saccade a decision network was used to predict the digit class of the currently observed stimulus. The decision network received the activity of the recurrent layer as input. It’s prediction was used as input to the latent network together with *h*_*t*_ to produce samples *z* from *p*(*z* | *x, y*). These samples together with the class prediction ŷ were used as input to the decoder to predict the full image as samples *x*_*i*_ ∼ *p*(*x* | *z, y*).

The decision network was instantiated as another 2-layer MLP with leaky ReLU activation in the first layer and a softmax output in the final classification layer. The first layer consisted of 128 units and the last layer of 10 units for the 10 target classes. As the network’s classification was used as input to the latent style representation as well as to the decoder, the additional classification task had explicit influence on the prediction of the target image and, as a consequence, on the model uncertainty and saccade patterns.

In order to learn to correctly classify the digit labels, an additional loss term was added at the end of each saccade sequence. The classification loss was calculated by the cross-entropy, or negative log likelihood, between the predicted softmax probabilities *p*(ŷ | *x*) and the one-hot-encoded target class *y*

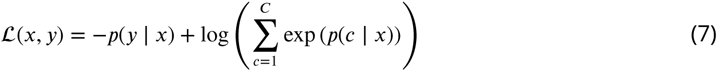

for *C* possible classes. The total resulting loss term form Experiment II was given by

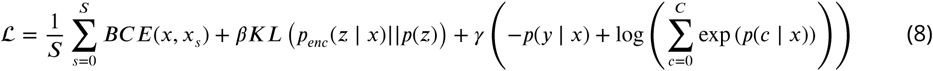

where *γ* was an additional weighting term for the relevance of the classification task. As for the weighting of the KL-divergence, the classification weight was slowly increased over the course of the training with *γ* = tanh(*v* · *epoch*) with hyperparameter *v. v* was set to 0.005 for all experiments.

## Acknowledgments

SQ and MG were supported by VIDI grant number 639.072.513 of The Netherlands Organization for Scientific Research (NWO).

